# Regulatory mechanisms for the axonal localization of tau in neurons

**DOI:** 10.1101/456608

**Authors:** Minori Iwata, Shoji Watanabe, Ayaka Yamane, Tomohiro Miyasaka, Hiroaki Misonou

## Abstract

Tau is a microtubule (MT)-associated protein, which precisely localizes to the axon of a mature neuron. Although it has been widely used as an axonal marker, the mechanisms for its axonal localization have been elusive. This might be largely due to the lack of an experimental system, as exogenously expressed tau, such as GFP-tau, mis-localizes to the soma and dendrites. In this study, we found that the expression of endogenous tau and its axonal localization in cultured rat hippocampal neurons mainly occur during early neuronal development and are coupled. By mimicking this early expression, we demonstrate that exogenously expressed human tau can be properly localized to the axon, thereby providing the first experimental model to study the mechanisms of tau localization. Using this model, we obtained surprising findings that the axonal localization of tau did not require the MT-binding domain nor correlate with the MT-binding ability. Instead, we identified a transport mechanism mediated by the proline-rich region 2 (PRR2), which contains a number of important phosphorylation sites. Mimicking phosphorylation and dephosphorylation in PRR2 disrupts the axonal localization, suggesting that it is regulated by the phosphorylation state of PRR2. These results shed new lights on the mechanism for the axonal localization of tau and indicate a link between the hyperphosphorylation and mis-localization of tau observed in tauopathies.

**Significance statement:** In this paper, we present a first experimental system, in which expressed tau is properly localized to the axon, and which can therefore be used to study the mechanisms of tau localization and mis-localization. Using this system, we provide evidence that the microtubule binding domain of tau nor stable binding of tau to microtubules is not necessary for its axonal localization. Instead, we identified the proline-rich region and its phosphorylation-state dictate the localization of tau in neurons. These results provide a novel foundation to consider how axonal tau mis-localize to the soma and dendrites during early pathogenesis of Alzheimer’s disease.

## INTRODUCTION

Tau is a microtubule (MT)-associated protein (MAP) localized in the axon of a neuron in normal physiological conditions (Binder et al., 1985; Peng et al., 1986; Brion et al., 1988; Migheli et al., 1988; Trojanowski et al., 1989). In contrast, in Alzheimer’s disease (AD) and related disorders, tau mis-localizes to the soma and dendrites and forms neurofibrillary tangles. Understanding this dramatic alteration is key to understand the pathogenesis of AD, as tau pathology is unambiguously linked to neurodegeneration from pathological and genetic evidence (Gómez-Isla et al., 1997; Delacourte et al., 1999; Ghetti et al., 2015). To do so, we need to understand how normal tau is localized to and maintained in the axon and how these mechanisms are impaired. Previous studies have made considerable efforts to study and understand the mechanisms for the axonal localization of tau. These studies have revealed essentially two major mechanisms. First, tau is transported to the axon by MT- and motor protein-based mechanisms (Utton et al., 2005; Falzone et al., 2009; Scholz and Mandelkow, 2014). Second, MT-based filter situated near or in the axon initial segment restricts tau from entering the soma while allowing it to go into the axon (Li et al., 2011).

However, one big challenge remaining in studying these mechanisms is to localize exogenous tau to the axon, where endogenous tau localizes exclusively (Mandell and Banker, 1996). When tau is expressed in cultured neurons by transfection, it typically distributes uniformly throughout the cells (Xia et al., 2016), such that the major population of tau remains in the soma and dendrites (see Fig. 1B). This ‘mis-localization’ makes it difficult to study axonal localization mechanisms of tau in a precise manner. We therefore set to tackle this challenge and establish a good experimental model for studying tau localization.

**Figure 1.**
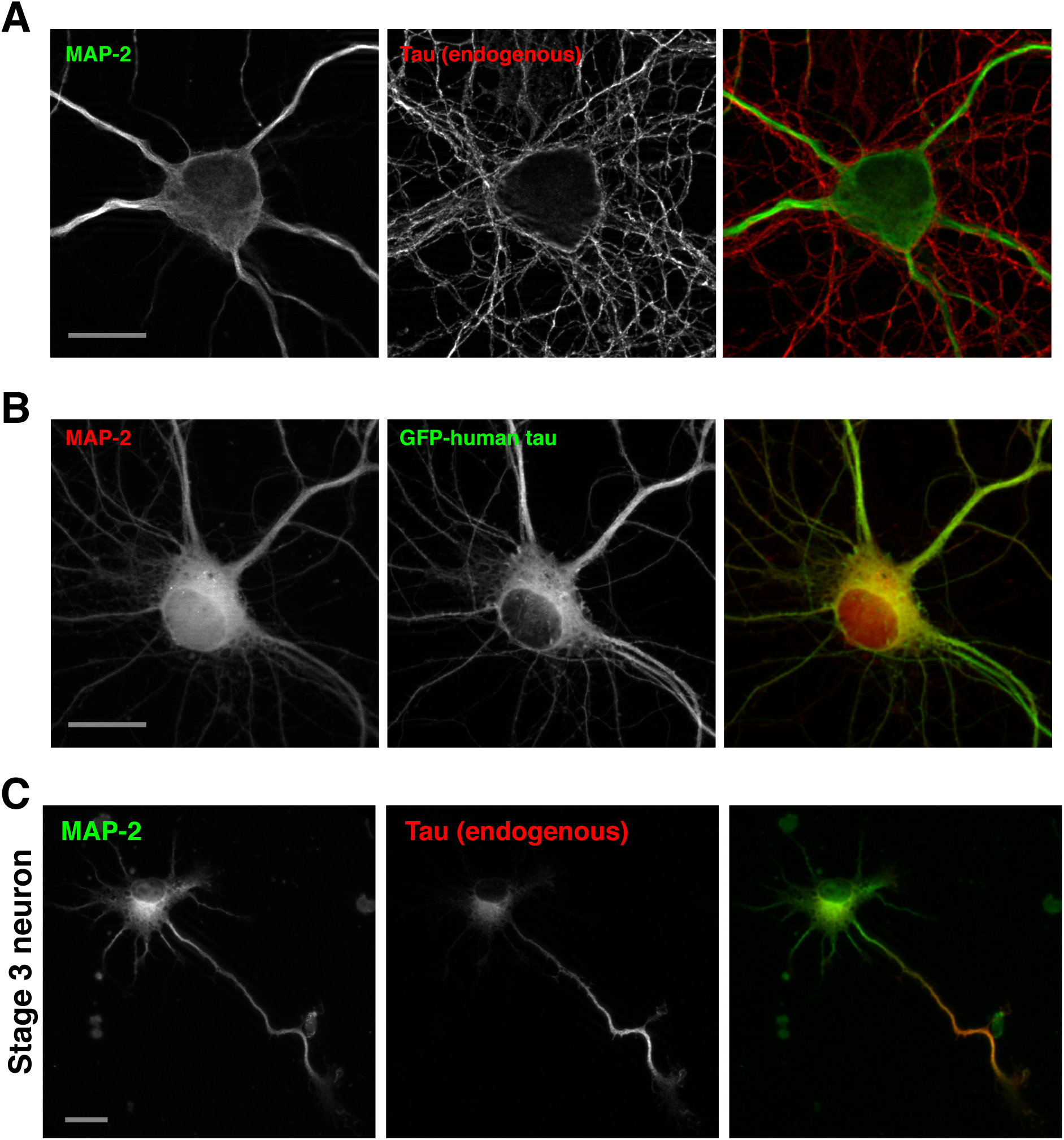
Axonal localization of endogenous tau and mis-localization of exogenously expressed tau. **A,** Axonal localization of endogenous tau in cultured neurons immunostained for MAP-2 (green) and tau (red). Scale bar, 20 *µ*m. **B,** Mis-localization of exogenous human tau tagged with GFP (green) overlapping with MAP-2 immunolabeling (red). Scale bar, 20 *µ*m. **C,** Early axonal localization of endogenous tau in stage 3 neurons at 2~3 DIV. It should be noted that at this stage MAP-2 expressed at low levels is uniformly distributed throughout neurons. Scale bar, 10 *µ*m.

In our accompanying paper, we showed that tau mRNA expression is high during prenatal to the first week of postnatal development in mice. We also demonstrated that expressing human tau in the same manner as endogenous tau results in its axonal localization like endogenous tau in knock-in mice, whereas human tau expressed beyond the developmental period mis-localizes to the soma and dendrites in tau transgenic mice. In this paper, we show that this holds true in cultured neurons, such that exogenous human tau can be localized to the axon only when it is expressed early during neuronal development. Using an inducible expression system, we were able to establish an experimental model in which exogenous tau is colocalize with endogenous tau in the axon in both developing and matured neurons. With this model, we show novel findings on the relationship between MT-binding and axonal localization of tau, the involvement of the proline-rich region in tau localization, and how phosphorylation affects the localization and MT-binding.

## MATERIALS AND METHODS

### Neuronal culture

Dissociated cultures of embryonic (E17~18) rat hippocampal neurons were prepared from female timed pregnant rats as previously described (Misonou et al., 2008) with minor modifications. Briefly, dissected hippocampi were digested in 0.25% trypsin for 15 min at 37°C, dissociated by pipetting, and then plated onto glass coverslips coated with 1 mg/ml poly-Llysine at 50,000 cells/coverslip. Neurons were cultured for 3 to 14 days in vitro (DIV) in 6-well plates, on the bottom of which contained astrocyte cultures. Coverslips were lifted with wax pedestals as described by Kaech and Banker (Kaech and Banker, 2006). Cytosine arabinoflanoside was added to the culture at 2 DIV to prevent the growth of non-neuronal cells on the coverslips. All animal use was approved by the institutional animal care and use committee.

### DNA constructs

Human tau cDNA corresponding to the 0N4R variant (1 - 363 amino acid residues) was amplified and cloned into pRK172 using NdeI and EcoRI sites. For the deletion mutants, PRR2 and MTBD, corresponding regions (140 - 185 amino acid residues for PRR2 and 186 - 309 for MTBD) were removed using QuickChange mutagenesis. The coding regions of tau wild-type and mutants were amplified using PCR and cloned into pAcGFP (Clontech). Synthesized phospho-mimetic and dephospho-mimetic fragments (eurofins Genomics) were introduced into SacII and PstI site of pAcGFP-Tau. Resultant AcGFP-tau and mutants (GFP-tau hereafter) was then amplified using PCR and cloned into pLVSIN lentiviral vector (Clontech), of which CMV promoter was replaced with the TRE3GS element/promoter from pTetONE (Clontech).

### Lentiviral vectors

The production of lentiviral particles was carried out as described previously (Chen and Okayama, 1987). HEK Lenti-X 293T cells were maintained in Dulbecco’s Modified Eagle Medium (Nacalai Tesque) with 10 % Fetal Bovine Serum (SIGMA), 1 *µ*g/ml penicillin and streptomycin in 5% CO_2_ at 37°C until use. Cells were re-plated at 1 x 10^6^ cells per 10 cm culture dish before a day of transfection. Lentiviral plasmids (6 *µ*g of pRSV-Rev, pMD2.G, and pMDLg/p RRE plasmids from Addgene #12253, 12259, and 12251, respectively) were mixed with 12 *µ*g of pLVSIN containing GFP-tau and 12 *µ*g of pRRLSIN.cPPT.PGK-GFP.WPRE (Addgene #12252), of which EGFP was replaced with Tet3G (Clontech). The mixture was then mixed with 50 *µ*l of 2.5 M CaCl_2_, and supplemented with sterilized water to adjust the volume to 500 *µ*l. They were incubated for 20 min at room temperature (~25°C) after being mixed with 2x BES-buffered saline (50 mM BES sodium salt, 280 mM NaCl, 1.5 mM Na_2_HPO_4_, pH6.95). Transfection mixture was added onto cells, which were then transferred to a 3% CO_2_ incubator. Next day (after about 16-20 h), the culture medium was exchanged with 10 ml of neuronal culture medium. Transfected cells were transferred to 5% CO_2_ incubator and incubated for 2 days. Culture medium was collected and filtered through a 0.45 *µ*m filter unit into a 15 or 50 ml conical tube. Collected lentiviral solution was kept at 4°C for short-term preservation or at − 80°C for long-term storage. For lenti virus titer measurement, p-24 lenti virus titer kit (TaKaRa) was used.

### Inducible expression of tau

At the day of plating, hippocampal neurons were infected with the lentiviral solution at 10 ~ 50 MOI overnight. Briefly, coverslips with neurons were transferred to new 6-well plates with 1~1.5 ml of astrocyte conditioned media in each well. The conditioned media were prepared as described previously (Misonou and Trimmer, 2005). After the overnight incubation, the media were replaced with fresh conditioned media with 1 *µ*g/ml doxycycline for expression induction. Coverslips were transferred back to the original 6-well plates with astrocytes after 1 h induction.

### Immunofluorescence labeling

Neurons were fixed in 4 % Paraformaldehyde/PBS for 20 min. Blocking and permeabilization was done in 0.2% fish skin gelatin/0.1 % Triton X-100/TBS. Primary and secondary antibodies were diluted in the same buffer and applied in separate incubation steps of 1 h and 45 min, respectively. Coverslips were mounted on glass microscope slides using ProLong Gold anti-fade reagent (Thermo Fischer Scientific). We used the following primary antibodies (and dilutions): rabbit anti-rodent tauN raised against the peptide (DTMEDHAGDYTLLQDEG) corresponding to the N-terminal potion of mouse tau (serum at 1:1,000), rat anti-total tau (RTM38) raised against purified recombinant human tau (),

### Localization analysis

The steady-state localization of tau and other marker proteins was documented using a Olympus IX73 microscope with a 60x/1.42 NA objective lens and Andor Zyla5.5 camera, or a Carl Zeiss ApoTome system with a 63x/1.4 NA lens. Line scan analysis of fluorescence intensities was performed using ImageJ.

### Fluorescence recovery after photobleaching (FRAP)

Neurons expressing GFP-tagged proteins were imaged using an Olympus FV-1000 microscope equipped or Carl Zeiss LSM 700. Images were taken every 1 s. Photobleaching was induced by applying the 450 nm laser to a circular (3.5 *µ*m diameter) or a rectangular spot at 100% laser power for 200 ms. Fluorescence intensity was measured using ImageJ, corrected for background fluorescence and the overall bleaching due to imaging, and normalized for the maximum and minimum fluorescence intensities.

### Microtubule-binding assay

Recombinant tau proteins were purified from *Escherichia coli* (*E. coli*) transformed with pRK172 vectors encoding his-tagged human tau and deletion mutants, as previously described (Xie et al., 2014). The purity was verified with SDS-PAGE and Coomassie brilliant blue staining. Microtubules were prepared as described previously with minor modifications (Planel et al., 2007). Briefly, mouse brains were homogenized in MT stabilization buffer (1 mM MgSO_4_, 1 mM EGTA, 0.1 mM DTT, 0.5% Triton X100, 10% glycerol, 100 *µ*M taxol, 2 mM GTP, protease/phosphatase inhibitors, and 0.1 M MES, pH 6.8). After a brief centrifugation to remove debris, the supernatant was then centrifuged at 135,000 x g for 15 min at 2°C. The resultant pellet was resuspended in the MT stabilization buffer containing 0.5 M NaCl to strip endogenous tau from microtubules and centrifuged again. The pellet was resuspended in the MT stabilization buffer, and this MT fraction was used for the binding assay immediately.

Purified tau protein (250 nM as the final concentration) and 100 *µ*l of the MT fraction were mixed and incubated on ice for 30 min. The mixture was centrifuged at 135,000 x g for 20 min at 2°C. The levels of tau in the supernatant (MT-unbound fraction) and in the pellet (MT-bound fraction) were quantified using Western blotting with purified recombinant tau as standards.

### Experimental Design and Statistical Analysis

All statistical analyses were performed on GraphPad Prism software (GraphPad Software, Inc. CA, USA). Power analysis was performed using G*Power (Faul et al., 2007, 2009) with parameters taken from previous reports or similar experiments.

FRAP data were background subtracted, compensated for bleaching from imaging, and normalized to make the minimum and maximum 0 and 1, respectively (Jensen et al., 2017). The recovery phase from the minimum were fitted with the one-phase association model: Y= Y_0_ + (Plateau − Y_0_) x (1 − 10^-k x X^). The difference of the fitted curves among different tau variants or in different areas in neurons were analyzed with F-test for the null-hypothesis that a curve with shared parameters (slope k and Plateau) fits better than those with different parameters for each data set. We also fitted data from individual neurons and obtained averaged slope and Plateau for each mutant. For Fig. 3B, the recovery data were fitted with a one-dimensional diffusion model (Ellenberg and Lippincott-Schwartz, 1999) to obtain the estimate of diffusion coefficient for WT tau.

For the analysis of axonal transport, intensity profiles over the bleached area from three neurons were obtained from right after photobleaching and 10 s (WT tau) or 5 s (ΔPRR2) later and averaged for each time point. The area between these two curves was then measured at proximal and distal areas within the bleached area and compared using paired Student’s t-test.

For the MT-binding data, we normalized MT-bound fractions to those of WT tau in each experiment to compensate for the variability in MT stability. The normalized data from all tau variants were analyzed at once using ANOVA with Dunnett’s post hoc tests to minimize multiple comparisons.

To compare the localization of tau mutants relative to that of endogenous tau, we measured average fluorescence intensities of expressed tau in a dendrite and the axon and computed the ratio of axonal over dendritic signals. The value was normalized to that of endogenous tau in the same neuron. We obtained normalized ratios from 6~8 neurons for each tau mutant from at least two independent cultures. The averages were compared to that of WT tau using ANOVA with Dunnett’s post hoc tests.

## RESULTS

### A new experimental system in which exogenous tau is localized to the axon

Tau is commonly used as an axonal marker because of its precise localization to the axon, as shown in Fig. 1A. To study the mechanism of this axonal localization, researchers have used cultured neurons to express exogenous tau with a tag, such as GFP. However, expressed tau is uniformly expressed in the soma, dendrites, and the axon (Fig. 1B), such that one cannot use this kind of experimental systems to study the precise localization mechanisms. We therefore sought to address this issue and to establish an experimental system in which exogenous tau localizes to the axon.

Typically, cultured neurons are transfected at 5 ~ 10 DIV, and tau expression is induced constitutively by a strong promoter such as a CMV promoter. However, it has been reported that endogenous tau is highly expressed and localized to the axons in immature neurons (Mandell and Banker, 1996). We confirmed that tau is preferentially localized to the axon even in the stage 3 neurons, which are right after the axon specification period (Fig. 1C). We also found that the relative intensity of tau immunoreactivity is high at 3 DIV and decline as neurons become mature (Fig. 2A). Based on these observations, we hypothesized that for exogenous tau to be properly localized to the axon it needs to be expressed transiently during early development (Fig. 2B).

**Figure 2.**
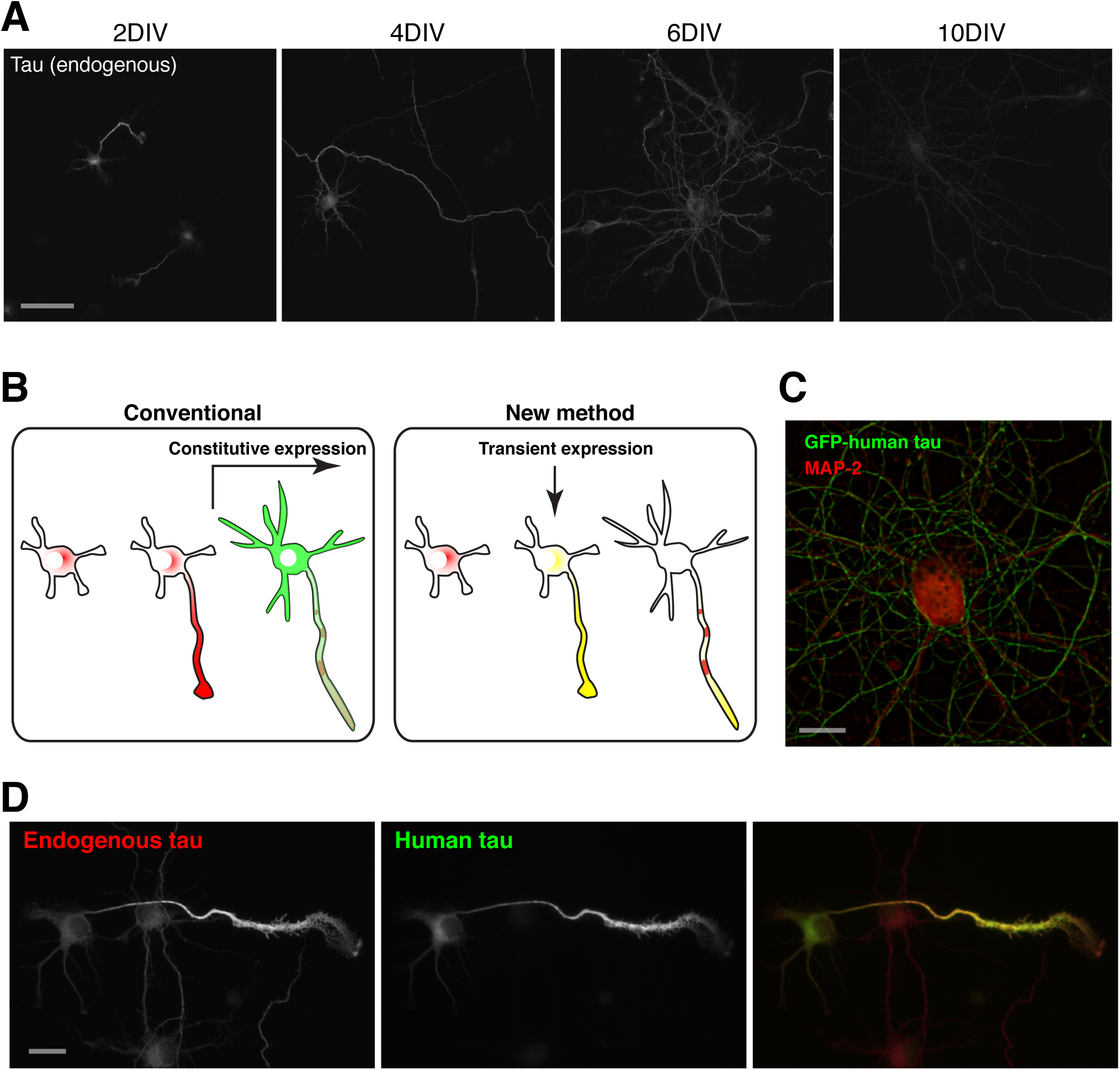
Normal axonal localization of exogenous tau by following the expression profile of endogenous tau. **A,** Developmental expression and localization of endogenous tau in cultured neurons. All images were taken at a fixed camera exposure. Scale bar, 50 *µ*m. **B,** Scheme illustrating our method. Red and green indicate endogenous and exogenous tau, respectively. Endogenous tau is expressed in stage 2 neurons and localized to the axon in stage 3 neurons. In conventional methods, exogenous tau is expressed constitutively beyond this stage and mis-localizes to the soma and dendrites. In our new method, the expression of exogenous tau is induced in immature neurons around stage 2 and 3 only transiently. This results in axonal localization of exogenous tau. **C,** Axonal localization of exogenous human-tau when expressed transiently in young neurons infected with GFP-tagged tau (green) and mKate2-tagged MAP-2C (red) at 1 DIV. It should be noted that there was virtually no overlap between GFP-tau and mKate2-MAP-2. Scale bar, 20 *µ*m. **D,** Reproduction of the early axonal localization of endogenous tau (red) with GFP-tagged tau (green). Scale bar, 20 *µ*m.

To test it, we constructed Tet-ON lentiviral vectors, with which the expression of exogenous tau can be controlled pharmacologically via the tetracycline transactivator (Urlinger et al., 2000). Cultured neurons were infected with lentiviral particles at 0 DIV and treated with doxycyclin to induce the expression for 1 h at 1 DIV. This transient expression resulted in axon specific localization of human tau (Fig. 2C). Although the expression was induced only for the short period of time (1 h), GFP-tau was readily detected even in the 14 DIV neurons, indicating that the turnover of tau is slow as previously implicated (Mercken et al., 1995; Morales-Corraliza et al., 2009; Yamada et al., 2015; Sato et al., 2018). Furthermore, we were able to recapitulate the developmental localization of tau. As shown earlier in Fig. 1C, endogenous tau is concentrated to the distal axon even in stage 3 neurons. GFP-tau expressed under the Tet-ON system also showed this preferential localization to the axon and co-localization with endogenous tau (Fig. 2D).

We then tested if it is the timing of expression or the briefness of the expression by expressing GFP-tau transiently at 7 DIV. Fig. 3A shows the axonal localization of GFP-tau in 14 DIV neurons, in which the expression was induced at 1 DIV. Line scan analysis indicates that GFP-tau is co-localize with endogenous tau in the axon but not with MAP-2 in and dendrites (Fig. 3B). In contrast, GFP-tau was readily detected in the soma with MAP-2, when expressed at 7 DIV (Fig. 3C). Line scan analysis also revealed that GFP-tau was mostly in dendrites rather than the axon in these neurons (Fig. 3D). These results support our hypothesis and demonstrate a novel experimental system to study the mechanisms for the axonal localization of tau, and presumably for its mis-localization.

**Figure 3.**
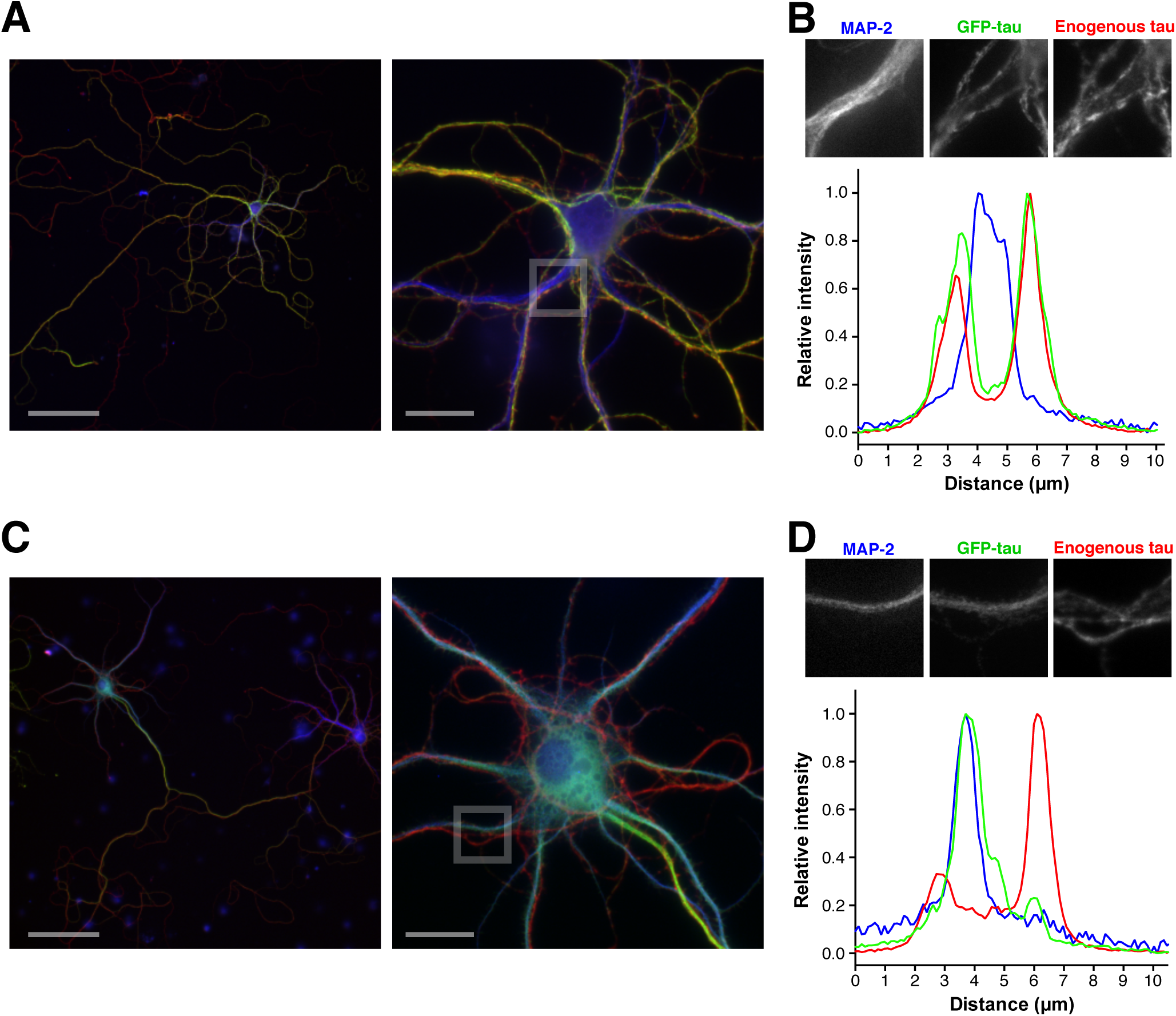
Axonal localization of tau dependent on the timing of expression. **A,** Triple immunolabeling of neurons, in which the expression of WT tau was induced at 1 DIV, for WT tau, endogenous tau, and MAP-2 at 14 DIV. Both low (left panel) and high (right panel) magnification images are shown. Scale bars, 100 *µ*m (left) and 20 *µ*m (right). **B,** Line scan analysis of endogenous tau (red), WT tau (green), and MAP-2 (blue) in the neuron shown in A. Top panels show high magnification images of the area indicated in the right panel in A, which were used for the analysis. **C,** Triple immunolabeling of neurons, in which the expression of WT tau was induced at 7 DIV, for WT tau, endogenous tau, and MAP-2 at 14 DIV. Both low (left panel) and high (right panel) magnification images are shown. Scale bars, 100 *µ*m (left) and 20 *µ*m (right). **D,** Line scan analysis of endogenous tau (red), WT tau (green), and MAP-2 (blue) in the neuron shown in C. Top panels show high magnification images of the area indicated in the right panel in C, which were used for the analysis.

### Molecular mobility of exogenous tau unraveling its MT-binding

Diffusional motion and MT-binding of tau have been studied in neurons using fluorescence recovery after photobleaching (FRAP) and single-molecule tracking techniques (Konzack et al., 2007; Weissmann et al., 2009; Janning et al., 2014). Here we also tested how exogenous tau behaves in our experimental system using FRAP. We observed somewhat slow mobility of GFP-tau in distal axons in neurons at 3 DIV (Fig. 4A and 4B) with the half time of recovery at about 15 s. We fitted the recovery phase with a one-dimensional diffusion model (Ellenberg and Lippincott-Schwartz, 1999) and obtained the diffusion coefficient of 0.15 ± 0.01 *µ*m^2^/s (Fig. 4B), which is much smaller than that previously reported. This slow mobility in the axon persisted during the maturation of neurons, such that there were only small differences in FRAP at 3, 7, and 14 DIV (Fig. 4C). In contrast, the mobility of GFP-tau in the soma changed substantially and was significantly decreased between 3 and 14 DIV (Fig. 4D and 4E). These results indicate that GFP-tau binds to MTs more stably in the axon than previously reported, and that MT-binding of tau is facilitated in the soma.

**Figure 4.**
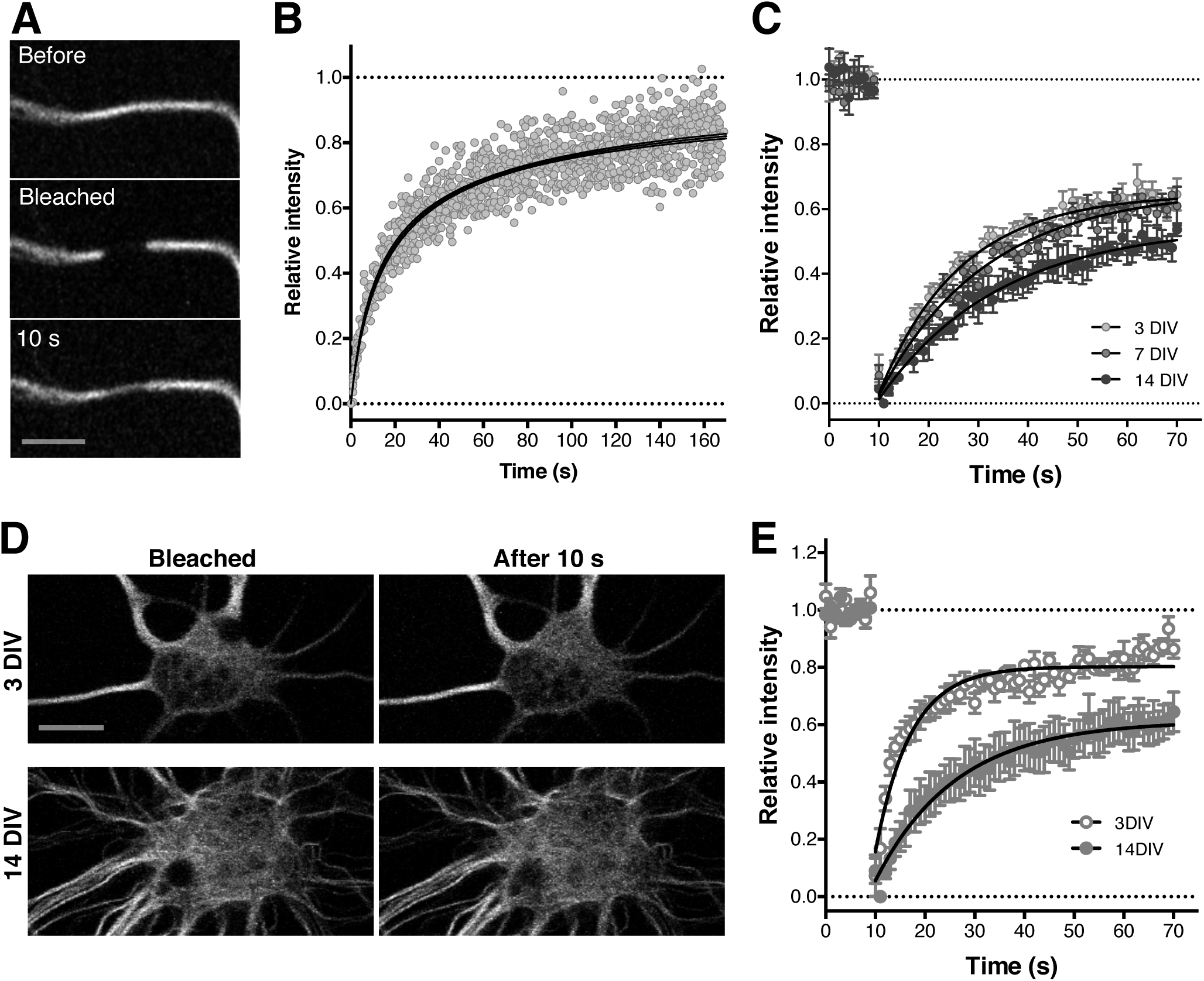
Diffusional properties of tau in neurons. **A,** FRAP of GFP-tagged human tau in the axons of cultured neurons at 3 DIV. FRAP was performed in a middle portion of the axon. Images before, immediately after bleaching, and 10 s after bleaching are shown. Scale bar, 5 *µ*m. **B,** Recovery rate of fluorescence from four neurons. Individual data points are shown with grey circles. The data were fitted with a one-dimensional diffusion model with the diffusion coefficient of 0.15 ± 0.01 *µ*m^2^/s, which is shown as the solid line with 99% confidential interval (dotted lines). **C,** Recovery rate of fluorescence in the axon from neurons at 3, 7, and 14 DIV. The data shown are the mean ± S.E.M and were fitted with exponential functions (solid lines). **D,** FRAP of GFP-tagged tau in the somata of cultured neurons at 14 DIV. Images immediately after bleaching and 10 s after bleaching are shown. Scale bar, 10 *µ*m. **E,** Recovery rate of fluorescence in the somata. It should be noted that the somatic signals were very weak compared to those in the axon albeit detectable. The data shown are the mean ± S.E.M. and were fitted with exponential functions. The slopes are significantly different between 3 and 14 DIV (p< 0.0001 with F (1, 594) = 32.21 using regression analysis with exponential functions).

To test if the mobility of GFP-tau reflects its MT-binding *in situ*, we generated tau lacking the MT-binding domain (MTBD) (Fig. 5A) and investigated its MT-binding in an *in vitro* assay (Fig. 5B) and in FRAP in neurons. As expected, recombinant tau lacking MTBD (tau ΔMTBD) purified from E. Coli exhibited significantly reduced MT-binding (p< 0.0001 with q (30) = 7.145 using ANOVA with Dunnett’s test, analysis done altogether with all the mutants as shown in Table 1), whereas wild-type tau (WT tau) was detected mostly in the MT-bound fraction (Fig. 5C, and see Table 1 for details). We then performed FRAP to probe their MT binding in the axons of cultured neurons. As shown in Fig. 5D, in contrast to the slow recovery of wild-type tau, GFP-tagged ΔMTBD showed a virtually complete and faster recovery. When the normalized fluorescence signals were fitted with single exponential functions (Fig. 5E), the slopes and the immobile fraction differ significantly between WT tau and tau ΔMTBD (Slope, 0.062 ± 0.004 vs 0.267 ± 0.024; immobile fraction, 0.37 ± 0.02 vs 0.12 ± 0.03, both p< 0.0001) (see Table 2 for details). Regression analysis also showed a significant difference in slope (p< 0.0001 with F (1, 715) = 295.4). These characteristics (p= 0.75 for WT tau 3 vs 7 DIV, and p= 0.10 for ΔMTBD using ANOVA with Sidak’s post hoc tests) and the difference (p< 0.0001, WT tau vs ΔMTBD at 7 DIV with regression analysis) were maintained in more mature neurons at 7 DIV for both WT tau and ΔMTBD. These results suggest that WT tau binds to MTs in the axon and that that ΔMTBD diffuses freely and does not efficiently bind to microtubules.

**Figure 5.**
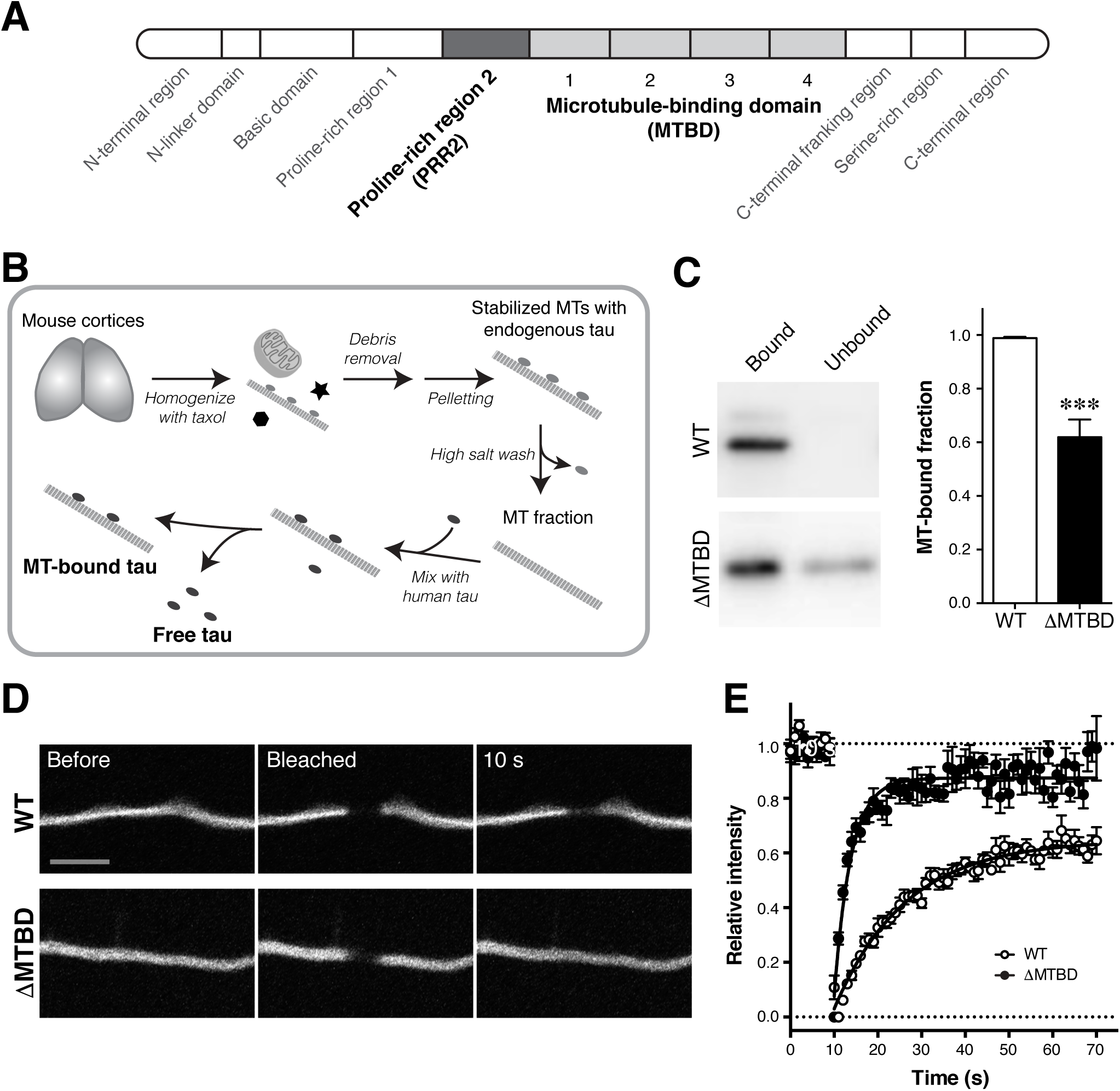
Microtubule-binding of tau. **A,** Schematic diagram illustrating structural and functional domain of human tau. **B,** Biochemical assay of the microtubule (MT)-binding of tau *in vitro*. Briefly, mouse cortices are homogenized in a warm buffer with taxol to stabilize microtubules. MTs are obtained by high speed centrifugation from the post-nuclear supernatant. Bound mouse tau is removed by washing MTs with 0.5 M NaCl. This microtubule fraction is mixed with recombinant human tau for the binding assay. **C,** MT binding of wild-type (WT) tau and tau lacking the MT-binding domain (ΔMTBD) in vitro. WT tau and ΔMTBD in MT-bound and -unbound fractions were measured using quantitative Western blotting (left panels). The data were normalized to the values of WT tau and shown as the mean ± S.E.M (p< 0.0001 ANOVA with Dunnett’s multiple comparison test). **D,** FRAP of WT tau and ΔMTBD in the axons of cultured neurons at 3 DIV. FRAP was performed in a middle portion of the axon. Images before, immediately after bleaching, and 10 s after bleaching are shown. Scale bar, 5 *µ*m. **E,** Recovery rate of fluorescence. The recovery of ΔMTBD was significantly faster than that of WT tau (p< 0.0001).

**Table 1.** Microtubule-binding of tau mutants.

|  | **MT-bound** | **MT-unbound** |
| --- | --- | --- |
| **WT tau** | 0.989 ± 0.005 | 0.011 ± 0.005 |
| **ΔMTBD\*\*\*** | 0.619 ± 0.066 | 0.381 ± 0.066 |
| **ΔPRR2\*** | 0.839 ± 0.012 | 0.161 ± 0.013 |
| **ΔPRR2-MTBD\*\*\*** | 0.059 ± 0.017 | 0.941 ± 0.017 |
| **PRR2_Glu\*\*** | 0.770 ± 0.062 | 0.230 ± 0.062 |
| **PRR2_Ala** | 0.993 ± 0.003 | 0.007 ± 0.003 |

Comparisons were performed using ANOVA with Dunnett's multiple comparison test. The overall difference was statistically significant ( $p < 0.0001$ with $F(5, 30) = 90.75$ . \* $p = 0.0295$ , \*\* $p = 0.0005$ , \*\*\* $p < 0.0001$ ).

**Table 2.**
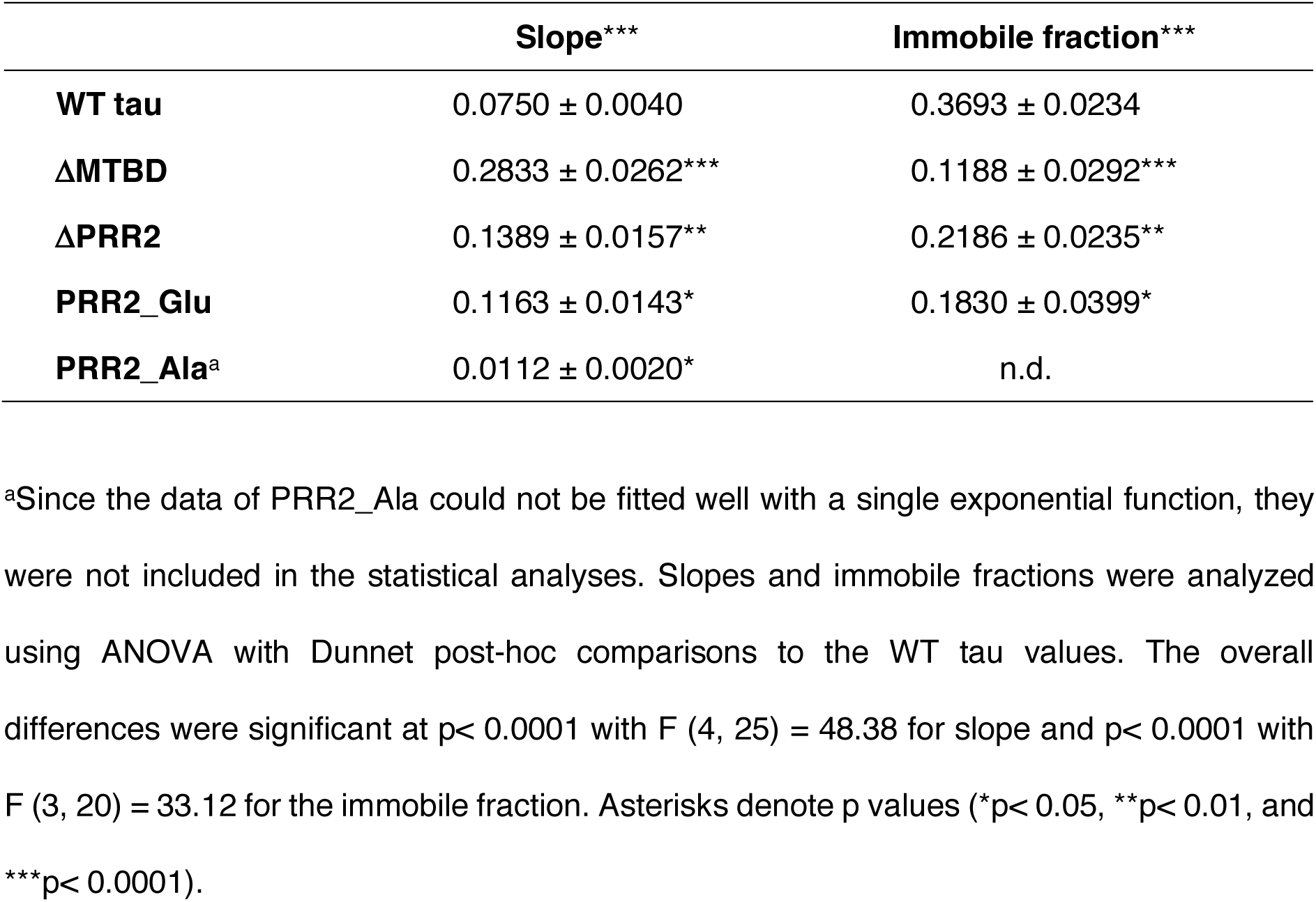
Curve fitting parameters using exponential functions for the FRAP data.

### Axonal localization of GFP-tau independently of the microtubule-binding

While we perform FRAP, we realized that ΔMTBD localized to the axon like WT tau in immature neurons at 3 DIV (Fig. 6A) and more mature neurons at 7DIV (Fig. 6B). This was very surprising because of the following reasons. First, it has been thought that the axonal localization of tau is achieved by its preferential binding to axonal MTs to dendritic MTs (Kanai and Hirokawa, 1995; Weissmann et al., 2009). Second, MT-dependent transport systems have been also implicated (Utton et al., 2005; Falzone et al., 2009; Scholz and Mandelkow, 2014). Third, it has been proposed that a MT-based molecular filter prevents tau from reentering the soma from the axon and therefore maintains the axonal localization of tau (Li et al., 2011).

**Figure 6.**
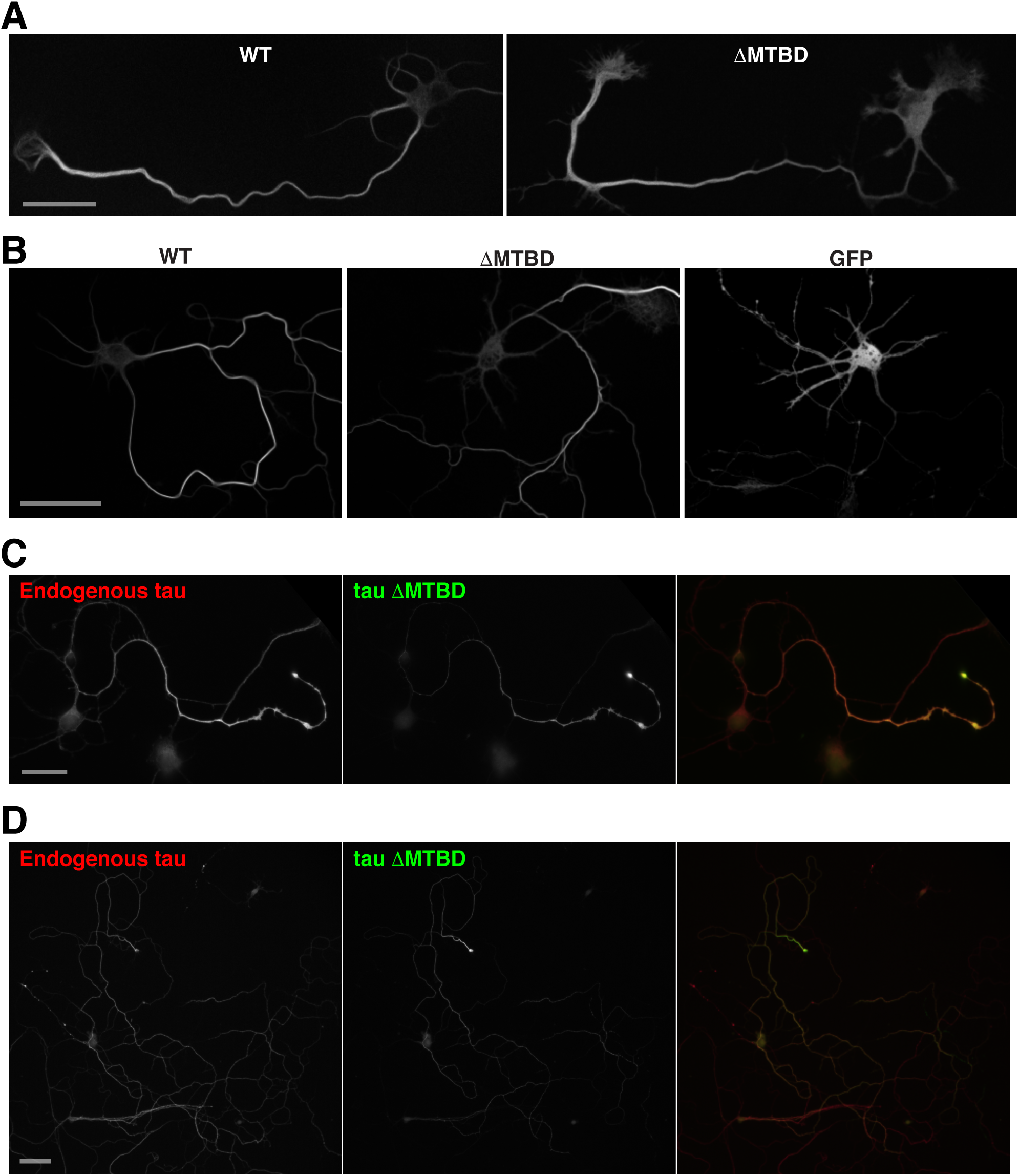
Axonal localization of tau independent of its microtubule-binding. **A,** Distribution of GFP-tagged WT tau and ΔMTBD in neurons at 3 DIV. Direct fluorescence signals from them are shown. Scale bar, 20 *µ*m. **B,** Distribution of GFP-tagged WT tau, ΔMTBD, and GFP alone in neurons at 7 DIV. Direct fluorescence signals from them are shown. It should be noted that dendritic and axonal development of neurons expressing WT tau and ΔMTBD appears normal. Scale bar, 50 *µ*m. **C,** Immunofluorescence labeling of endogenous tau (red) and ΔMTBD (green) in neurons at 3 DIV. Scale bar, 20 *µ*m. **D,** Immunofluorescence labeling of endogenous tau (red) and ΔMTBD (green) in neurons at 14 DIV. Scale bar, 50 *µ*m.

To further investigate the localization of ΔMTBD, we performed double immunolabeling with endogenous tau. It also revealed that ΔMTBD co-localizes with endogenous tau in the axon at 3 DIV (Fig. 6C). Since the axonal localization was evident with the direct GFP signals (Fig. 6A and 6B), the apparent axonal localization is not due to preferential labeling of axonal tau molecules in immunocytochemistry. It retained the axonal localization even in more mature neurons at 14 DIV (Fig. 6D). The axonal localization is also not a general tendency for any cytosolic proteins, as mKate2-tagged MAP-2 (Fig. 2C) and GFP (Fig. 5B) did not exhibit the axonal accumulation like WT tau and ΔMTBD. Therefore, these results suggest that the preferential localization of tau to the axon does not require MTBD and stable MT-binding.

### Axonal localization mediated by the proline-rich region 2 (PRR2)

Since the deletion of MTBD did not affect the localization of tau, we aimed to determine the region of tau determining its axonal localization using a series of deletion mutants. We found that removing the proline-rich region 2 (ΔPRR2) (see Fig. 5A) resulted in a uniform distribution of tau in the soma, dendrites, and the axon of neurons (Fig. 7A), which is strikingly different from WT tau and ΔMTBD (Fig. 6A). This mis-localization of ΔPRR2 is not due to its increased MT-binding. The biochemical MT-binding assay revealed that ΔPRR2 exhibits a significant albeit slight loss of MT-binding (p= 0.0295 with q (30) = 2.887) (Fig. 7C and Table 1). Interestingly, deletion of both PRR2 and MTBD resulted in a greater reduction of MT-binding than ΔMTBD and almost complete loss of MT-binding (p<0.0001 with q (30) = 17.95 against WT) (Fig. 7C and Table 1). FRAP analyses of ΔPRR2 also showed significant increases in the recovery rate (0.075 ± 0.004 vs 0.1389 ± 0.0157, p= 0.0087) and the mobile fraction (0.6307 ± 0.0234 vs 0.7814 ± 0.0235, p= 0.0011) compared to WT tau (Fig. 7B and Table 2), suggesting that PPR2 participates in MT-binding as previously indicated (Butner and Kirschner, 1991; Gustke et al., 1994; Goode et al., 1997).

**Figure 7.**
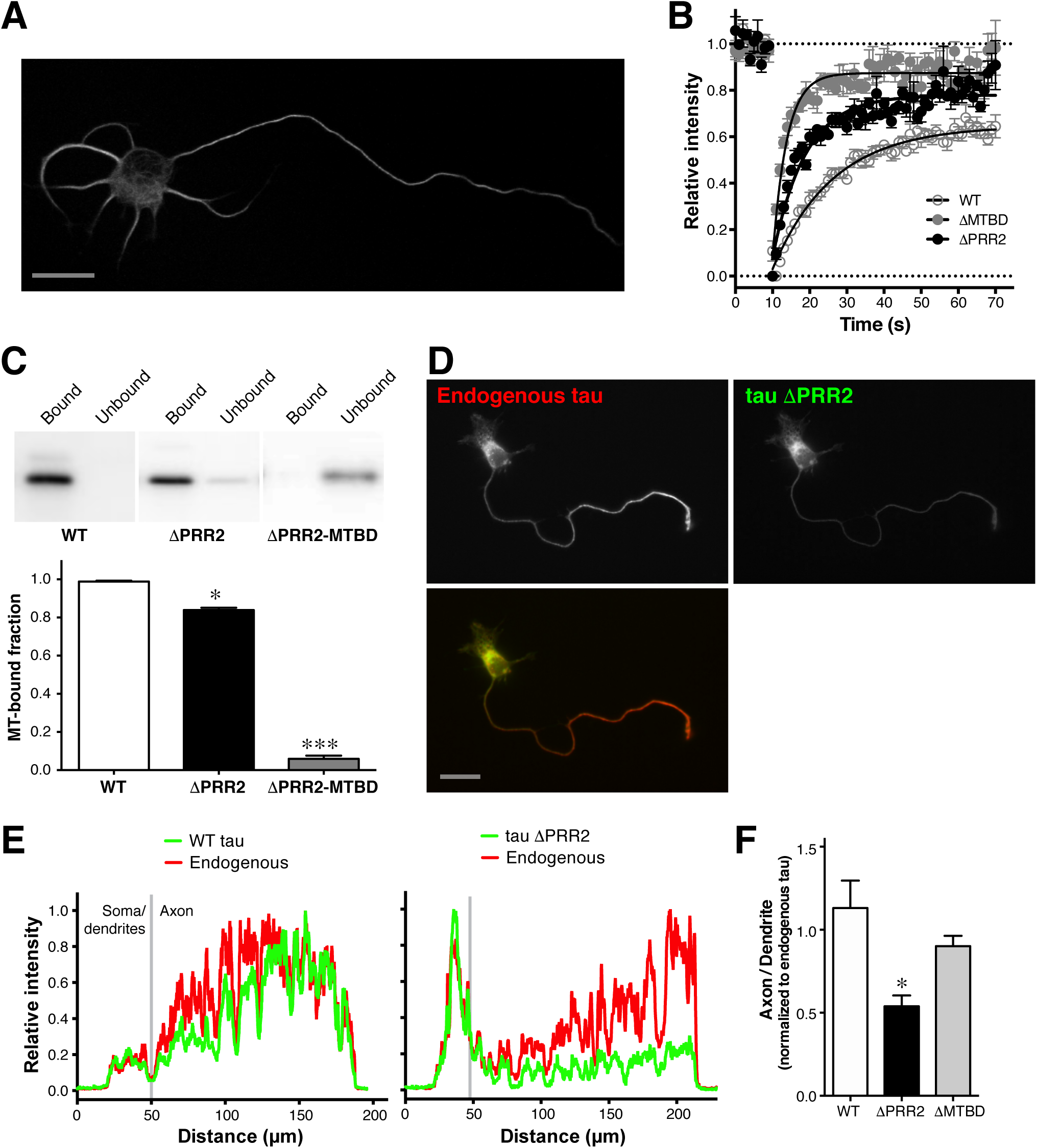
Axonal localization dependent of the proline-rich region 2. **A,** Distribution of GFP-tagged tau lacking the proline-rich region 2 (ΔPRR2) in neurons at 3 DIV. Direct fluorescence signals are shown. Scale bar, 20 *µ*m. **B,** FRAP of ΔPPR2 in the axons of 3 DIV neurons. Data from WT tau and ΔMTBD in Fig. 4D are also shown for comparison (p< 0.0001 vs WT tau with F (1, 723) = 34.87 using regression analysis). **C,** MT binding of WT tau, ΔPRR2, and tau lacking both PRR2 and MTBD (ΔPRR2-MTBD) in vitro. Tau in MT-bound and -unbound fractions was measured using quantitative Western blotting (left panels). *p= 0.0295. **D,** Immunofluorescence labeling of endogenous tau (red) and ΔPRR2 (green) in neurons at 3 DIV. Scale bar, 20 *µ*m. **E,** Line scan analysis of endogenous tau (red) and either WT tau or ΔPRR2 (green). The vertical grey lines indicate the border between the soma and the axon. **F,** Quantification of how exogenous tau is enriched in the axon like endogenous tau using the ratio of axonal signals over dendritic signals normalized to those of endogenous tau. Note that a good overlap of them would provide a value close to 1. *p= 0.0199 with q (28)= 2.989 using ANOVA and Dunnett’s post hoc test.

To further quantify the effect of PRR2 deletion on the localization, we performed double-immunolabeling of ΔPRR2 and endogenous tau. As shown in Fig. 7D, ΔPRR2 localization was dramatically different from that of endogenous tau. This difference was also evidenced by the line scan analysis of fluorescence signals (Fig. 7E). We also computed the ratios of axonal signals over dendritic signals for endogenous and exogenous tau (ΔMTBD and ΔPRR2) and normalized the ratio of exogenous tau to that of endogenous tau in each neuron. Therefore, a good colocalization of exogenous tau with endogenous tau would result in a value close to one. As shown in Fig. 7F, WT tau, which exhibits a similar enrichment in the axon as endogenous tau (Fig. 2D), had the ratio close to one. However, ΔPPR2 tau showed a significantly smaller ratio (0.9069 ± 0.1478 vs 0.4851 ± 0.0675, p= 0.0199, q (28) = 2.989, ANOVA with Dunnett’s post hoc test, analysis done altogether with all the mutants as shown in Fig. 11G and Table 3), showing that it is not enriched in the axon. In contrast, there was no difference between WT and ΔMTBD (p= 0.6788). This mis-localization was maintained in more mature neurons as shown in Fig. 7. While WT tau was only detected in the axon with endogenous tau (Fig. 8A and 8B), ΔPPR2 was present in the soma and dendrites as well as the axon (Fig. 8C and 8D). These results suggest that PRR2 is critical for the axonal localization of tau.

**Table 3.** Localization of tau mutants

|  |  |
| --- | --- |
| **WT tau** | 0.9069 ± 0.1478 |
| **ΔMTBD** | 0.7644 ± 0.0805 |
| **ΔPRR2** | 0.4851 ± 0.0675* |
| **PRR2_Glu** | 0.5035 ± 0.0841* |
| **PRR2_Ala** | 0.5028 ± 0.0770* |

Comparisons were done using ANOVA with Dunnett's post hoc tests. The overall difference was significant ( $p=0.013$ with $F(4, 28) = 3.864$ ). Pairwise comparisons with WT tau were performed using Dunnett's test (\* $p < 0.05$ ).

**Figure 8.**
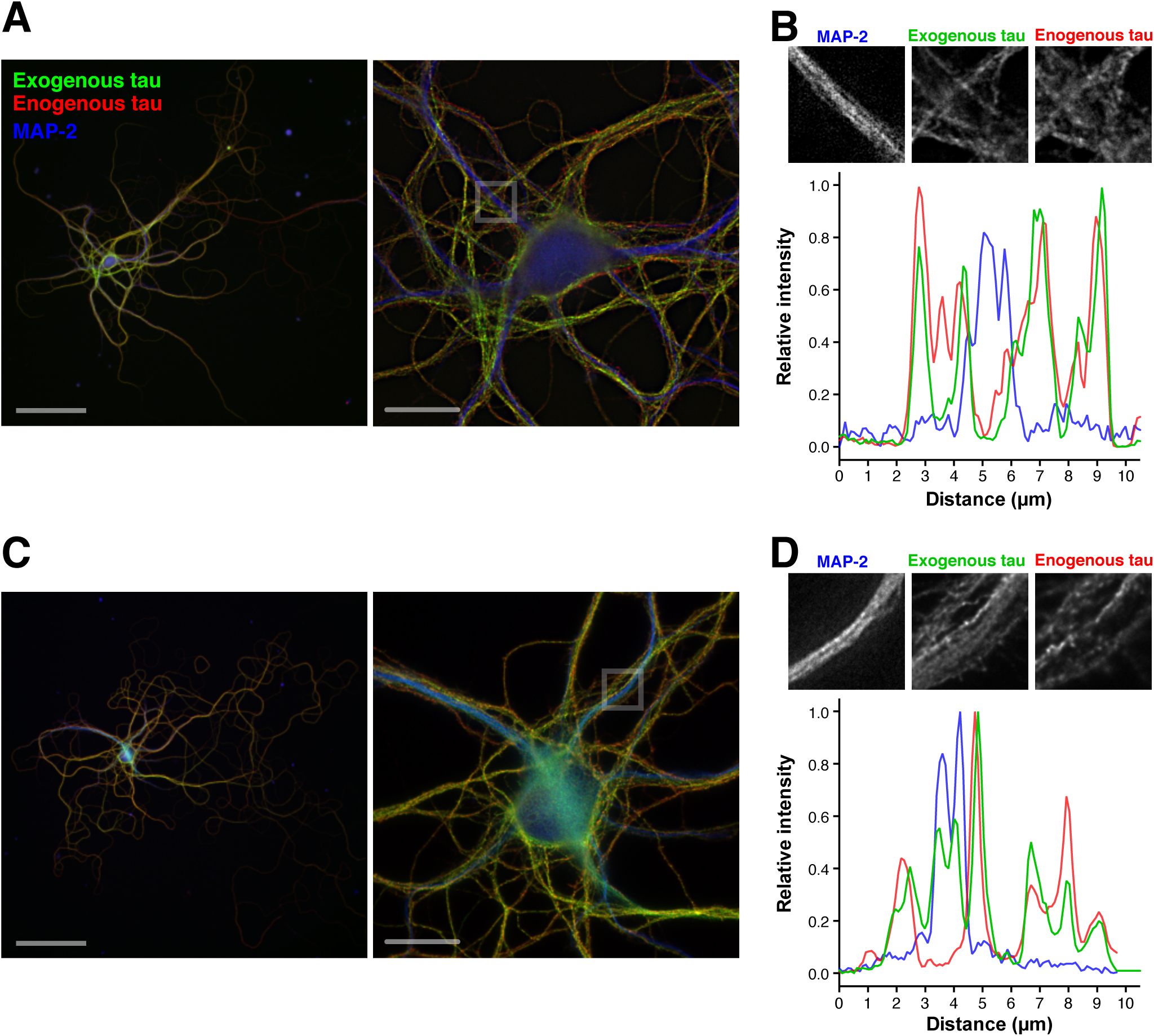
Mis-localization of ΔPPR2 in mature neurons. **A,** Triple immunolabeling of WT tau, endogenous tau, and MAP-2 in a 24 DIV neuron. Both low (left panel) and high (right panel) magnification images are shown. Scale bars, 100 *µ*m (left) and 20 *µ*m (right). **B,** Line scan analysis of endogenous tau (red), WT tau (green), and MAP-2 (blue) in the neuron shown in A. Top panels show high magnification images of the area indicated in the right panel in A, which were used for the analysis. **C,** Triple immunolabeling of ΔPPR2, endogenous tau, and MAP-2 in a 24 DIV neuron. Both low (left panel) and high (right panel) magnification images are shown. It should be noted that there are substantial fluorescence signals of ΔPPR2 (green) in the soma. Scale bars, 100 *µ*m (left) and 20 *µ*m (right). **D,** Line scan analysis of endogenous tau (red), ΔPPR2 (green), and MAP-2 (blue) in the neuron shown in C. Top panels show high magnification images of the area indicated in the right panel in C, which were used for the analysis.

### Active transport of tau to the axon

We initially considered three plausible mechanisms for tau to localize to the axon (Fig. 9A). One obvious mechanism was that axonal MTs traps tau in the axon. However, we reject this model because ΔMTBD was diffusible anywhere in neurons even near the tip of the axon, where it is highly accumulated (Fig. 9B and 9C). The diffusivity was also comparable in dendrites and the axon (Fig. 9D). Also, its high diffusivity indicates that ΔMTBD does not bind to any large structures, such as actin filaments and neurofilaments. Therefore, we sought to test the alternative models.

**Figure 9.**
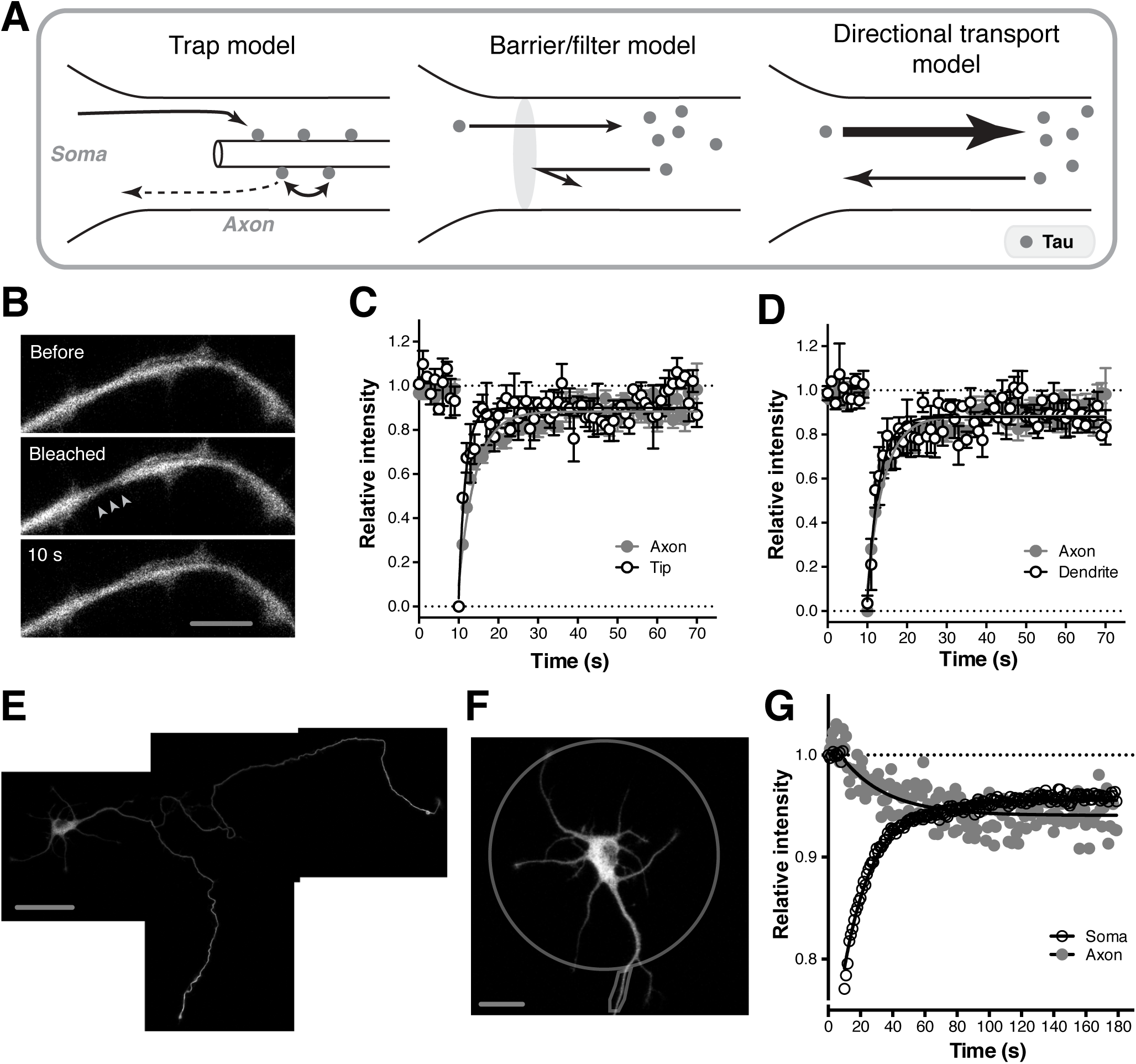
Potential mechanisms for the axonal localization of tau. **A,** Potential models for the axonal localization of tau. The trap model assumes a stable scaffold like microtubules, which binds and traps tau (grey circles) in the axon. In the barrier/filter model, a putative barrier/filter prevents tau molecules (or complexes) from escaping from the axon, while it allows somatic tau to enter the axon. The directional transport model proposes that an active transport mechanism constitutively sends tau from the soma to the axon at a rate greater than that of tau being sent bay by diffusion. **B,** FRAP of WT tau near the tip of the axon. Images before, immediately after bleaching, and 10 s after bleaching are shown. Scale bar, **C,** Rate of recovery from the experiment in B. The data shown are the mean ± S.E.M. and those in Fig. 3E is also shown for comparison. Curve fitting was done with an exponential function. The recovery was significantly faster in the tip than in the shaft of the axon (p< 0.0001 with F (1, 541) = 33.46 using regression analysis and F-test). This is presumably due to that the tips are typically thicker than the shaft. **D,** FRAP of WT tau in dendrites and axons. The data shown are the mean ± S.E.M. and those in Fig. 3E is also shown for comparison. Curve fitting was done with an exponential function. There was a significant difference in the rate of recovery between dendrites and axons (p= 0.006 with F (1, 601) = 7.57 using regression analysis and F-test). This is also probably due to the larger volumes of dendrites than axons. **E,** Neuron expressing ΔMTBD at 3 DIV. Scale bar, 50 *µ*m. **F,** Region of the neuron in E used for FRAP. The large circle indicates the bleached region, and the small eclipse shows the adjacent non-bleached area in the axon used to measure the concomitant decrease of fluorescence. Scale bar, 20 *µ*m. **G,** Changes of fluorescence during FRAP shown in F. Open circles show the recovery in the somatodendritic region. Grey closed circles show the reduction of fluorescence in the adjacent region. It should be noted that signals were not scaled to make the minimum values zero, unlike the other FRAP plots.

It has been proposed that the axon initial segment functions as a size filter for cytoplasmic proteins and organelles. Mandelkow and colleagues have also proposed that there is a filter or barrier which allows the entrance of tau into the axon but prevent it from returning to the soma (Li et al., 2011). To test If there is a barrier/filter like this, we performed FRAP of ΔMTBD in the soma. Neurons expressing ΔMTBD at 3DIV (Fig. 9E) were subjected to somatic FRAP. We expected that there was no significant recovery of ΔMTBD in the soma, as axonal ΔMTBD would not re-enter the soma. However, when the entire somatodendritic region was photobleached as shown in Fig. 6F, there was a rapid and substantial recovery of the somatodendritic fluorescence as well as a concomitant decrease of the axonal fluorescence (Fig. 9G). This suggests that tau ΔMTBD is freely diffusible from the axon to the soma and dendrites, in spite of its overall axonal localization. These results do not support the barrier/filter model.

We next sought to examine the directional transport of tau to the axon using FRAP. Although it has been previously proposed that tau is transported by a motor-MT transport system in the axon, there must be a MT-independent transport of tau to the axon. To do so, GFP-tau was photobleached in the beginning part of the axon as shown in Fig. 10A. This allowed us to monitor the flow of fluorescent molecules from proximal and distal sides and to test if there is a directional motion of tau. We found that WT tau exhibited an asymmetric flow after photobleaching, such that the fluorescence signal recovered preferentially from the somatic side of the bleached area (Fig. 10A). In contrast, ΔPRR2 exhibited more symmetric recovery (Fig. 10A).

**Figure 10.**
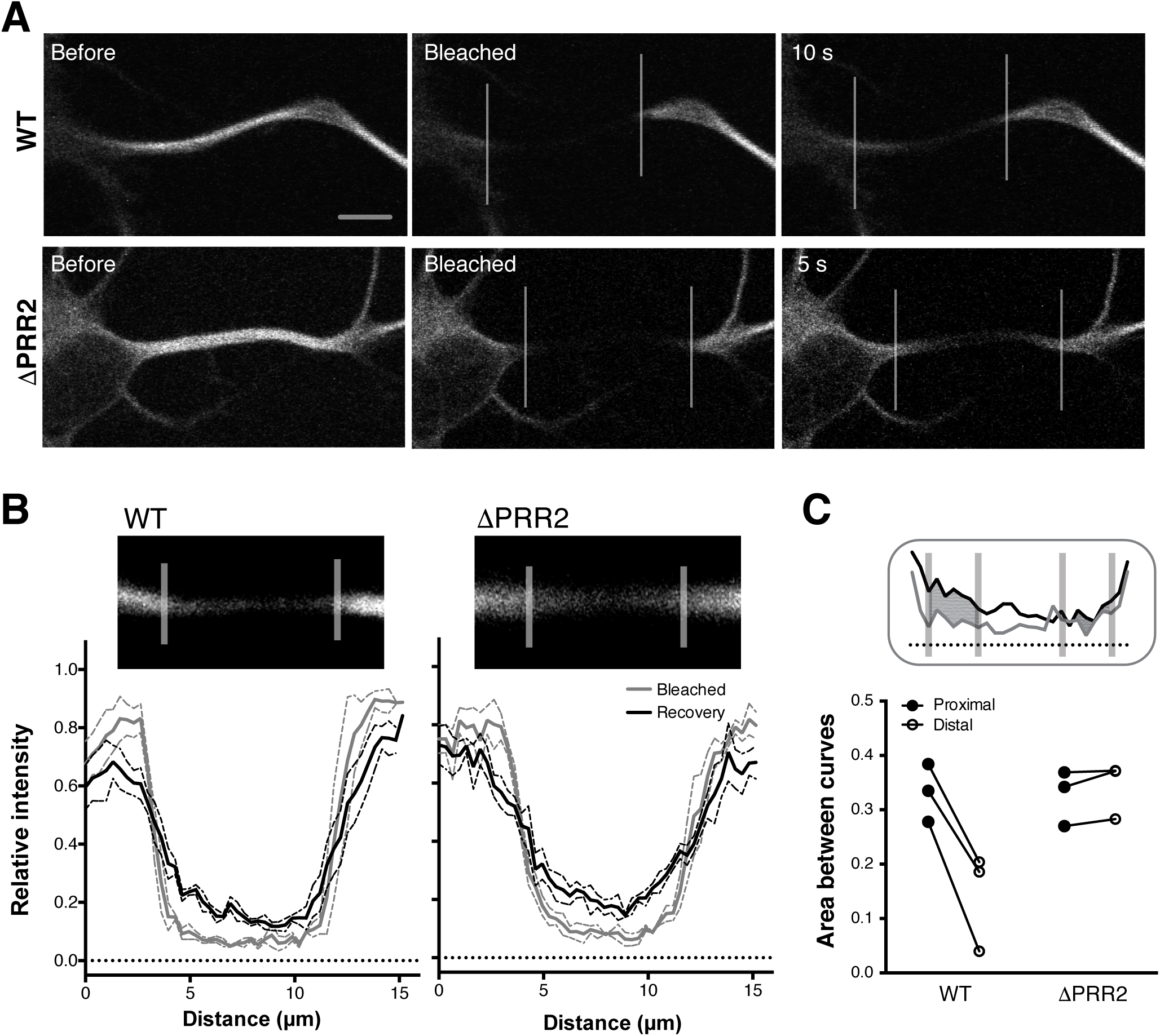
Directional transport of tau to the axon mediated by the PRR2 domain. **A,** FRAP in large areas in the proximal axon in neurons expressing WT tau or ΔPRR2. Vertical lines indicate the bleached regions. Images before, immediately after bleaching, and 10 s (WT) or 5 s (ΔPRR2) after bleaching are shown. The earlier time point was chosen for ΔPRR2 because of its faster diffusion than WT. Scale bar, 5 *µ*m. **B,** Spatial patterns of recovery. Fluorescence intensity on a line drawn over the bleached area was measured immediately and 5 or 10 s after bleaching and plotted against distance. Mean values from three neurons were plotted with S.E.M. (dotted lines). **C,** Fraction recovered was measured as the area between curves near the proximal and distal sides in the bleached area as illustrated in the inset. There is a significant difference between the proximal and distal values of WT (p= 0.0012 with t (4) = 9.841 using repeated measures two-way ANOVA with Sidak’s post hoc tests) but not ΔPRR2 (p= 0.7280).

To better quantify this difference, we performed FRAP in the axon where the thickness and fluorescence signals are uniform (Fig. 10B). The recovery was monitored using a line profile function and plotted against the distance. As shown in the line graphs, the recovery was asymmetric for WT tau but not for ΔPRR2. We measured the area under the curve in the proximal and distal bleached areas as indicated in Fig. 10C. This showed that the recovery was significantly greater (p= 0.0012 with t (4) = 9.841 using repeated measures two-way ANOVA with Sidak’s post hoc tests, n= 3) in the proximal side of the bleached area than the distal side for WT tau, whereas no significant difference was observed for ΔPRR2 (p= 0.7280). Taken together, these results suggest that ΔPRR2 is necessary for the axonal localization of tau and the directional transport of tau to the axon.

### Phosphorylation of PRR2 on MT-binding and axonal localization

PRR2 contains several phosphorylation sites, which are known to be hyper-phosphorylated in neurons in AD patients. Based on our results that PRR2 is important for the axonal localization of tau, and because tau is mis-localized in affected neurons in AD, we hypothesized that the phosphorylation states of PRR2 impact the localization of tau. To gain insight on this, we first generated tau, of which all 8 putative phosphorylation sites in PRR2 (Fig. 11A) are replaced with either alanine (Tau PRR2_Ala) to mimic dephosphorylation or glutamate (tau PRR2_Glu) as pseudo-phosphorylation. In the biochemical assay, PRR2_Glu showed a small decrease in MT-binding (p= 0.0005 with q (30) = 4.503), whereas PRR2_Ala was found virtually only in the bound fraction and indistinguishable from WT (Fig. 11B). They also showed very different characteristics in FRAP. Tau PRR2_Glu exhibited faster recovery compared to WT tau in axons, which is similar to ΔPRR2 (Fig. 11C). Surprisingly, tau PRR2_Ala showed a dramatic reduction in FRAP with a significantly reduced slope compared to WT (p= 0.013), which indicates its stable binding to MT *in situ*. Since the deletion of PRR2 did not abolish MT-binding per se, these results suggest that phosphorylation of PRR2 strongly regulates MT-binding of tau in neurons.

**Figure 11.**
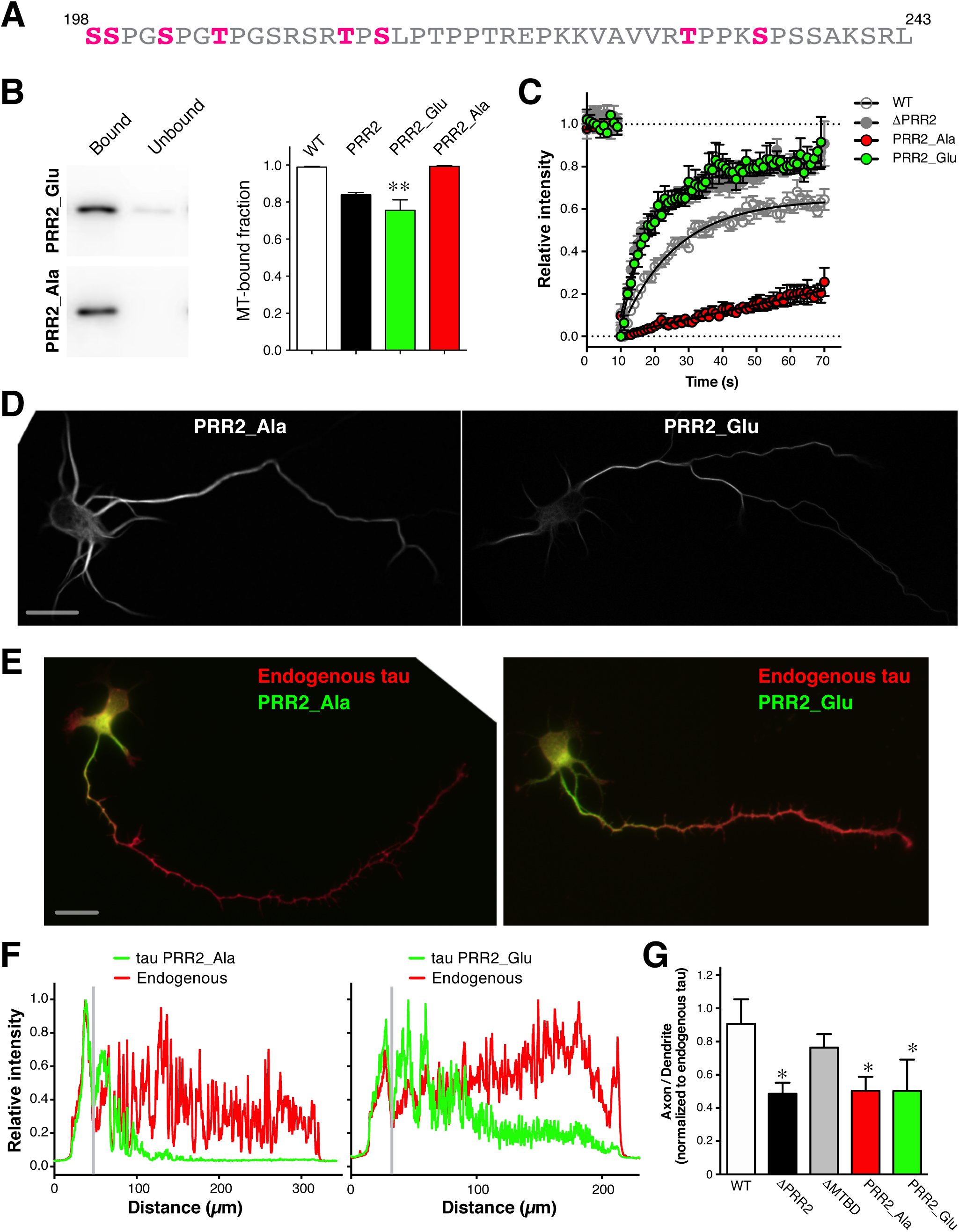
Regulation of tau localization via phosphorylation of PRR2. **A,** Amino acid sequence of PRR2. The bold letters indicate the eight putative phosphorylation sites mutated to Ala (PRR2_Ala) or Glu (PRR2_Glu). **B,** Microtubule-binding of PRR2_Ala and PRR2_Glu *in vitro*. The data of ΔPPR2 are also shown for comparison. **p= 0.0005. **C,** FRAP of PRR2_Ala and PRR2_Glu in the axon of neurons at 3 DIV shown in D. Data shown are the mean ± S.E.M. and fitted with exponential functions. Data of WT and ΔPRR2 in Fig. 6B are shown in grey symbols. **D,** Direct fluorescence images of PRR2_Ala and PRR2_Glu. Scale bar, 20 *µ*m. **E,** Immunofluorescence labeling of endogenous tau (red) and PRR2_Ala or PRR2_Glu (green) in neurons at 3 DIV. Scale bar, 20 *µ*m. **F,** Line scan analysis of fluorescence for endogenous and PRR2_Ala or PRR2_Glu in neurons shown in E. **G,** Ratios of axonal signals over dendritic signals normalized to those of endogenous tau. Data of WT and all mutants are shown. Comparison was done using ANOVA with Dunnett’s post hoc tests. *p< 0.05.

These mutants both exhibited abnormal axonal localizations. PRR2_Ala was highly accumulated in the soma, unlike WT tau and ΔPRR2 (Fig. 11D), which was also apparent with the double immunolabeling with endogenous tau (Fig. 11E). Line scan analysis clearly revealed that PRR2_Ala is stuck in the soma and proximal dendrites and cannot travel further (Fig. 11F), presumably due to its stable binding to MTs. PRR2_Glu was similar with ΔPRR2 in localization as shown in Figs. 11D, 11E, and 11F, such that it failed to accumulate in the distal axon. It also failed to show the asymmetric recovery in in the axon in FRAP (data not shown) as ΔPRR2 did. We also compared the axon/dendrite signal ratio normalized to that of endogenous tau. Both PRR2_Ala and PRR2_Glu had significantly lower ratio (0.5035 ± 0.0841, p= 0.021; 0.5028 ± 0.0770, p= 0.0268, respectively) than WT tau and comparable ratio to that of ΔPRR2 (Fig. 11G and Table 3). Therefore, hyper-phosphorylation could disrupt the function of PRR2 and PRR2-dependent axonal localization of tau, whereas hypophosphorylation enhances the MT-binding and also interfere the axonal localization.

## DISCUSSION

In this paper, we demonstrated a new cellular model to study the mechanisms for the axonal localization of tau. Our experimental model is unique in that exogenously expressed tau, such as GFP-tagged human tau, can be highly localized to the axon and colocalize with endogenous tau. This is different from previously used culture models, in which exogenous tau is constitutively expressed and distributes throughout the cell uniformly. Therefore, we believe that our experimental system provides a new platform to study the mechanisms of tau localization and mis-localization.

In combination with FRAP, our experimental model also allowed us to study MT-binding of tau in the axon *in situ*. The MT-binding of tau mutants assessed by FRAP was in a good agreement with that obtained in the in vitro biochemical assay. One potential caveat in the FRAP assay would be that excess amount of exogenous tau saturates MTs and results in an augmented pool of free tau. Our results indicate that this is the case with our system. First, the diffusivity of WT tau estimated from our FRAP analysis was smaller than that expected when a large portion of tau molecules is free. Second, PRR2_Ala mutant exhibited a negligible level of freely diffusible tau, while we observed comparable levels of direct GFP fluorescence from this mutant and all other tau constructs. Also, expression of WT tau did not affect the axonal localization of endogenous tau. These results suggest that expressed tau does not saturate MTs in our experimental model.

With the *in situ* MT-binding assay, we confirmed that both MTBD and PRR2 participate in MT-binding with MTBD being as the primary binding site (Goode et al., 1997). The deletion of PRR2 showed a modest effect on MT-binding in vitro and *in situ*. However, PRR2_Ala exhibited a significant increase in MT-binding in the *in situ* assay, indicating that the phosphorylation state of PRR2 impacts MT-binding, as previously reported (Kiris et al., 2011; Schwalbe et al., 2015). This effect was not observed in the biochemical assay because virtually all WT tau was already found in the MT-bound fraction, such that the binding could not be further improved in this assay. Recombinant WT tau, which is not phosphorylated, may just behave exactly like PRR2_Ala in vitro. In contrast, WT tau and PRR2-Ala were completely different from one another in FRAP. This suggests that WT tau in neurons is phosphorylated to some extent and exhibits more dynamic binding to MTs. This is consistent with that embryonic tau is highly phosphorylated in vivo.

PRR2_Ala also mis-localized to the soma and dendrites robustly. This suggests that tight and stable binding of dephosphorylated tau prevents it from going far in the axon. With that WT tau in neurons has reduced MT-binding, phosphorylation of PRR2 might be necessary for the initial transport of tau to the distal axons. Although we have not been able to identify the exact phosphorylation site(s) in PRR2 responsible for MT-binding thus far, this phosphorylation of PRR2 might be the reason why our method works to localize tau to the axon. It has been shown that tau is not highly phosphorylated in mature neurons in vivo in normal conditions, whereas it is in immature neurons. Therefore, tau expressed during the developmental stages is properly phosphorylated on select sites in PRR2, not tightly bound to MTs, and efficiently transported to the axon. In fact, our data showed that WT tau in immature neurons are significantly more mobile in the soma than in mature neurons. If tau is expressed beyond the developmental period or in mature neurons, it might be dephosphorylated, tightly binds to MTs, and therefore mis-localizes to the soma and dendrites.

Surprisingly, the axonal localization of tau was not dependent on its MT-binding via MTBD. ΔMTBD was localized to the axon with endogenous tau, despite that it did not seem to bind to MTs or any cytoskeletal structures in cells, evidenced by its rapid diffusional characteristics and the negligible level of immobile fraction in FRAP. In contrast, ΔPRR2 retained MT-binding but mis-localized to the soma and dendrites. Our FRAP data also indicated that the preferential transport from proximal to distal axon observed with WT tau was lost with ΔPRR2. Given that a large portion of molecules are readily diffusible for WT tau and ΔMTBD, these results suggest that there is a strong directional transport mechanism for tau, which can work against diffusional flux. In fact, we observed such directional transport of WT, which was lost with the PRR2 deletion.

It has been proposed that tau is transported to the axon by MT-dependent motors (Utton et al., 2005; Falzone et al., 2009; Scholz and Mandelkow, 2014). Our results show that MTBD is not necessary for the axonal localization of tau, although it is critical for the stable binding of tau to MTs in vitro and in the axon. These results indicate that MT-binding mediated by MTBD is not necessary for tau transport. However, the diffusivity of ΔMTBD in the axon we estimated was 2.25 ± 0.33 *µ*m^2^/s, which is still low for freely diffusible proteins. Therefore, it is possible that ΔMTBD binds transiently to MTs via PRR2 and/or linker protein(s), and is transported toward distal axons while repeating binding and unbinding with MTs. Our results that ΔMTBD retains MT-binding, presumably via PRR2, in the *in vitro* assay support this idea. Alternatively, tau is transported by MT-independent mechanisms mediated by actin filaments.

It has also been proposed that a filter or barrier near the axon initial segment prevents tau from entering the soma from the axon, thereby maintaining its axonal localization. Our results with ΔMTBD indicate that, even when a large portion of tau re-enters the soma, the overall axonal localization can be achieved, presumably by the active transport in developing neurons. It is still possible that a barrier mechanism helps to maintain the axonal concentration of tau, particularly in mature neurons. However, based on our results of ΔMTBD in mature neurons, we have to conclude that this kind of mechanism is largely MTBD-independent and/or has only a minor role in establishing the axonal localization of tau.

By employing the novel experimental system, our study confirmed the importance of PRR2 and its phosphorylation in MT-binding (Kiris et al., 2011; Schwalbe et al., 2015) *in situ* and further demonstrated its involvement in the axonal localization of tau. The results with the phosphorylation- and dephosphorylation-mimetic mutants indicate that phosphorylation of certain sites in PRR2 impairs the transport of tau to the axon and results in similar mis-localization as the deletion of the entire PRR2. In contrast, dephosphorylation of PRR2 would greatly enhance the binding of tau to MTs in the soma, thereby slowing its transport to the axon. Therefore, for tau to be localized in the axon, it has to be properly phosphorylated in the soma to benefit from the transport mechanism without being stuck on the somatic MTs. Our results also suggest that tau expressed during the developmental period is localized to the axon, while that expressed in mature neurons mis-localizes both in culture and *in vivo* as demonstrated in the accompanying paper (Kubo et al.). Considering these findings, we speculate that tau in AD is either ectopically expressed in the adulthood or becomes in excess due to the reduction of axonal MTs, and therefore mis-localized to the soma and dendrites.

## Acknowledgement

We thank Ms. Hongsun Park for her initial characterization of the inducible expression system, Drs. Nobuyuki Nukina, Tomoyuki Yamanaka, Haruko Miyazaki, and Naoto Saitoh for confocal microscopy, Drs. Shigeo Takamori and Yoshiaki Egashira for their technical assistance in lentivirus preparation. This work was supported in part by the Grant-in-Aid for Scientific Research on Innovative Areas (Brain Protein Aging and Dementia Control) (to T.M.) from MEXT, Core-to-Core Program A Advanced Research Networks, the Strategic Research Foundation at Private Universities (S1201009), and the JSPS KAKENHI Grant Number 26640030 (A.K.) and 16K07006 (to H.M.).

